# Pseudotime estimation: deconfounding single cell time series

**DOI:** 10.1101/019588

**Authors:** John Reid, Lorenz Wernisch

## Abstract

Cross-sectional time series single cell data confound several sources of variation, with contributions from measurement noise, stochastic cell to cell variation and cell progression at different rates. Time series from single cell assays are particularly susceptible to confounding as the measurements are not averaged over populations of cells. When several genes are assayed in parallel these effects can be estimated and corrected for under certain smoothness assumptions on cell progression. We present a principled probabilistic model with a Bayesian inference scheme to analyse such data. We demonstrate our method’s utility on public microarray, nCounter and RNA-seq data sets from three organisms. Our method almost perfectly recovers withheld capture times in an *Arabidopsis* data set, it accurately estimates cell cycle peak times in a human prostate cancer cell line and it correctly identifies two precocious cells in a study of paracrine signalling in mouse dendritic cells. Furthermore, our method compares favourably with Monocle, a state-of-the-art technique. We also show using held out data that uncertainty in the temporal dimension is a common confounder and should be accounted for in analyses of cross-sectional time series.

## 1 Introduction

### 1.1 Cross-sectional time series

Many biological systems involve transitions between cellular states characterised by gene expression signatures. These systems are typically studied by assaying gene expression over a time course to investigate which genes regulate the transitions. An ideal study of such a system would track individual cells through the transitions between states. Studies of this form are termed *longitudinal*. However current medium and high-throughput assays used to measure gene expression destroy cells as part of the protocol. This results in *cross-sectional* data wherein each sample is taken from a different cell.

This study analyses the problem of variation in the temporal dimension: cells do not necessarily transition at a common rate between states. Even if several cells about to undergo a transition are synchronised by an external signal, when samples are taken at a later time point each cell may have reached a different point in the transition. This suggests a notion of pseudotime to model these systems. Pseudotime is a latent (unobserved) dimension which measures the cells’ progress through the transition. Pseudotime is related to but not necessarily the same as laboratory capture time.

Variation in the temporal dimension is a particular problem in cross-sectional studies as each sample must be assigned a pseudotime individually. In longitudinal studies information can be shared across measurements from the same cell at different times.

Inconsistency in the experimental protocol is another source of variation in the temporal dimension. It may not be physically possible to assay several cells at precisely the same time point. This leads naturally to the idea that the cells should be ordered by the pseudotime they were assayed.

### 1.2 Single cell assays

The exploration of cell-to-cell heterogeneity of expression levels has recently been made possible by single cell assays. Many authors have investigated various biological systems using medium throughput technologies such as qPCR [1–4] and nCounter [5, 6] or high throughput technologies such as RNA-seq [7–14]. These studies have shown that cellular heterogeneity is prevalent in many organisms and regulatory systems. The variation in gene expression underlying this cellular heterogeneity has been attributed to several causes both technical and biological [3, 8–10]. Whilst accounting for variation in expression levels, none of these studies investigated how much is attributable to uncertainty in the temporal dimension. Conversely, methods such as Monocle and Wanderlust (described below) have attempted to place cells in a pseudotemporal ordering but do not explicitly model variation in the data.

### 1.3 Dimension reduction

Analyses of medium and high-throughput expression assays often use dimension reduction techniques. Anywhere between forty and several tens of thousands of gene expression levels may have been measured in each sample. This high-dimensional data can often be better analysed after projection into a low (two or three) dimensional latent space. Often this projection results in a natural clustering of cells from different time points or of different cell types which can then be related to the biology of the system. Such clusterings may suggest hypotheses about likely transitions between clusters and their relationship in time.

Dimension reduction has a large literature and there are many available methods, here we give a few examples of some that have been used in single cell expression analyses.

Principal components analysis (PCA) is prevalent in analyses of expression data [7, 8, 11, 14]. PCA finds linear transformations of the data that preserve as much of the variance as possible. In one example typical of single cell transcriptomics, Guo et al. studied the development of the mouse blastocyst from the one cell stage to the 64 cell stage [1]. They projected their 48-dimensional qPCR data into two dimensions using PCA. Projection into these two dimensions clearly separated the three cell types present in the 64 cell stage.

Multi-dimensional scaling (MDS) is another popular dimension reduction technique. MDS aims to place each sample in a lower dimensional space such that distances between samples are conserved as much as possible. Kouno et al. used MDS to study the differentiation of THP-1 human myeloid monocytic leukemia cells into macrophages after stimulation with PMA [3]. Their primary MDS axis explained the temporal progression through the differentiation, their secondary MDS axis explained the early-response of the cells to the stimulation they had undergone.

Independent components analysis (ICA) projects high dimensional data into a latent space that maximises the statistical independence of the projected axes. Trapnell et al. used ICA to investigate the differentiation of primary human myoblasts [12]. The latent space serves as a first stage in their pseudotime estimation algorithm Monocle (see below).

Gaussian process latent variable models (GPLVMs) are a dimension reduction technique related to PCA. They can be seen as a non-linear extension [15] to a probabilistic interpretation of PCA [16]. Buettner et al. used GPLVMs to study the differentiation of cells in the mouse blastocyst [17, 18]. They used qPCR data from Guo et al. who had analysed the expression of 48 genes in cells spanning the 1- to 64-cell stages of blastocyst development [1]. Buettner et al. were able to uncover subpopulations of cells at the 16-cell stage, one stage earlier than Guo et al. had identified using PCA.

The latent space in all of the methods above is unstructured: there is no direct physical or biological interpretation of the space and the methods do not directly relate experimental covariates such as cell type or capture time to the space. The samples are placed in the space only to maximise some relevant statistic, although the analysis often reveals some additional structure. For example, one axis may coincide with the temporal dimension of the data, or cell types may be clearly separated. In these cases the structure has been inferred in an unsupervised manner. However there is no guarantee that the methods above will uncover any specific structure of interest, for example, a pseudotime ordering.

Here we propose to impose an a priori structure on the latent space. In the model presented in this paper the latent space is one-dimensional and the structure we impose on the space relates it to the temporal information of the cell capture times. That is the latent space represents the pseudotime.

### 1.4 Pseudotime estimation

A number of methods have been proposed to estimate pseudotimes in gene expression time series. Äijö et al. proposed a temporal scaling method DyNB to estimate pseudotimes [19]. DyNB shifts the observed time by a multiplicative factor representing speed of transition through the process. It is applicable to longitudinal rather than cross-sectional time series. Äijö et al. modelled RNA-seq count data from human Th17 cell differentiation using a negative binomial distribution with a time-varying mean. The time-varying mean was fit using a Gaussian process over the scaled pseudotime space. They compared this pseudotime based model favourably with a similar model that only used the capture time points.

Trapnell et al. have developed the Monocle pseudotime estimation algorithm [12]. Monocle is a two-stage procedure: first it uses the ICA dimension reduction algorithm to map gene expression data into a low-dimensional space; second it finds the minimal spanning tree over the samples’ locations in this space. This spanning tree is used to assign a pseudotime to each cell. Trapnell et al. show how Monocle can be used to identify pseudotemporal ordering, switch-like changes in expression, novel regulatory factors and sequential waves of gene regulation.

Wanderlust is a graph-based pseudotime estimation algorithm developed by Bendall et al. [20]. Wanderlust arranges the high-dimensional input data into a nearest neighbour graph wherein cells that have similar expression profiles are connected. Wanderlust then applies a repetitive randomised shortest path algorithm to assign an average pseudotime to each cell. Bendall et al. used Wanderlust to analyse human B cell lymphopoiesis.

Both of the Monocle and Wanderlust algorithms do not explicitly make a connection between the cell capture times and the estimated pseudotime. This has two effects: first in the inference of the pseudotime, nonsensical results are possible such as pseudotimes whose order is far from the capture times; second the estimated pseudotimes are not on the same scale as the capture times, they are quantified in arbitrary temporal units.

### 1.5 Gaussian processes

Gaussian processes (GPs) are Bayesian models that are well suited to model expression profiles and capture the uncertainty inherent in noisy data. Bayesian inference in GPs can be performed analytically and provides posterior mean estimates with a full covariance structure. A GP is parameterised by a mean and a covariance function. For more details, Rasmussen and Williams have published a comprehensive review [21].

GPs have been used extensively to model time series and other phenomena in biological systems: Stegle et al. designed a two-sample test for differential expression between time series using GPs [22]; Honkela et al. used GPs to model expression profiles of transcription factors in an ODE based model of gene regulation [23]; Äijö et al. used GP models of regulatory functions to infer gene networks [24]; and Kirk et al. used GPs to model time series in a multiple dataset integration method [25].

## 2 Methods

### 2.1 Data

Our method has been designed to analyse single cell data but there is no technical reason why each sample must be from a cell. The model itself and notion of pseudotime would suit many cross-sectional data sets. Indeed one of the data sets used in our results section is from whole leaf microarray assays. However for consistency of explanation we refer to each sample as a cell in this paper.

Our method works on data with a simple structure. First, it expects gene expression data on a logarithmic scale, such as Ct values from qPCR experiments or log transformed counts from RNA-seq experiments. Second, it requires a *capture time* for each cell. This specifies at which time point that cell was sampled.

Our notation for the data is: *G* is the number of genes assayed; *C* is the number of cells sampled; *x*_*g,c*_ is the expression level of gene *g* in cell *c* where 1 *≤ g ≤ G* and 1 *≤ c ≤ C*; the capture time of cell *c* is *k*_*c*_ where *k*_*c*_ *∈* {*κ*_1_, *κ*_2_,…, *κ*_*T*_}; and *T* is the number of distinct capture times.

#### 2.1.1 Cell size adjustment

Single cell expression measurements can often contain per-cell biases that present as differences in basal levels of transcription. These can be caused by biological effects such as cell size or technical effects such as lysis efficiency or sequencing depth. We use a technique based on the method proposed by Anders and Huber to account for these effects [26].

Given a subset of the genes 𝒢 ⊆ {1, *…, G*} and a subset of the cells 𝒞 ⊆ {1, *…, C*} we define the cell size for cell c ∈ 𝒞 as

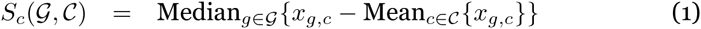

The median is used as it provides a robust estimate of the differences across genes. We estimate the cell sizes separately for the cells grouped by capture time,

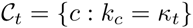

only using those genes that are expressed in at least half of the cells,

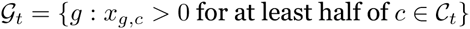

and apply the cell size as a correction to the raw data

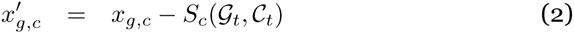

where *t* is such that *k*_*c*_ = *κ*_*t*_. In all that follows we use 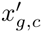 rather than *x*_*g,c*_ as our expression data.

### 2.2 Model

The primary latent variables in our model are the pseudotimes. The model assigns a pseudotime to each cell such that the induced gene expression profiles over the latent pseudotime space have low noise levels and are smooth.

Our model captures several aspects of the data: first, the data are noisy which we model in a gene-specific fashion; second, we expect the expression profiles to be smooth; third, we expect the pseudotime of each cell not to stray too far from its capture time.

The model can be split into several parts: one part represents the gene expression profiles; another part represents the pseudotimes associated with each cell; and another part links the expression data to the profiles.

#### 2.2.1 Gene expression profiles

The expression profiles are modelled using Gaussian processes. The expression profile *y*_*g*_ of gene *g* is a draw from a Gaussian process

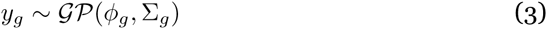

where *ϕ*_*g*_ is a (constant) gene-specific mean function estimated from the data and Σ_*g*_ is a gene-specific covariance function. The expression profiles are functions of pseudotime and as such the covariance function relates two pseudotimes.

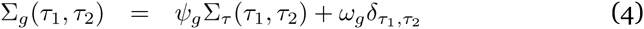

Here Σ_*τ*_ is a covariance function that defines the covariance structure over the pseudotimes. Σ_*τ*_ imposes the smoothness constraints that are shared across genes; *ψ*_*g*_ parameterises the amount of temporal variation this gene profile has; and *ω*_*g*_ models the noise levels for this gene. Log-normal priors for the *ψ*_*g*_ and *ω*_*g*_ are parameterised as

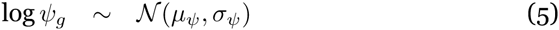

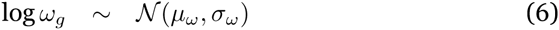

#### 2.2.2 Pseudotimes

The pseudotime *τ*_*c*_ for cell *c* is given a prior centred on the time the cell was captured.

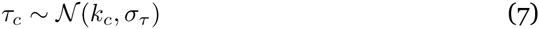

Each *τ*_*c*_ is used in the calculation of the covariance structure over pseudotimes Σ_*τ*_. Σ_*τ*_ is taken to be a Matern_3/2_ covariance function. Our experience shows that this function captures our smoothness constraints well although any reasonable covariance function could be used.

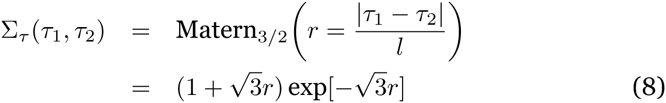

where *l* is a length-scale hyperparameter shared across the genes.

For cyclic data such as from the cell cycle or circadian rhythms we expect the expression profiles to be periodic. We can model this explicitly by a transformation of *r* in Equation (8). We replace *r* by *r*_Ω_

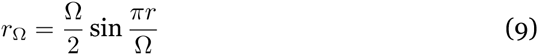

This has the effect of restricting the GP prior to periodic functions with period Ω.

#### 2.2.3 Expression data

The model links the expression data to the expression profiles by evaluating the profiles at the pseudotimes.

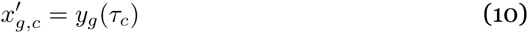

### 2.3 Relationship to other models

Briefly, our model can be interpreted as a one-dimensional GPLVM with a prior structure on the latent pseudotime space. The GPLVM model is a non-linear version of probabilistic PCA. In probabilistic PCA, the locations of the data in the latent space are given a Gaussian prior with zero mean and unit covariance. In our model the analogous latent variables are the pseudotimes. Our model gives the pseudotimes a structured prior rather than a standard normal: that is we relate the latent pseudotimes to the capture times of the cells using a Gaussian prior.

### 2.4 Hyperparameter estimation

All of the hyperparameters *μ*_*ψ*_, *σ*_*ψ*_, *μ*_*ω*_, *σ*_*ω*_ are estimated by an empirical Bayes procedure described below. The hyperparameters *l, σ_τ_* are supplied directly by the user of our method.

As with many hierarchical models, the parameters can have several posterior modes. For instance, much of the variation in typical single cell assay data could be explained by smooth expression profiles with high noise levels. Alternatively the same data could also be explained by rough expression profiles with low noise levels. Our model aims to balance these conflicting explanations and find parameters to fit the data with reasonable noise levels and expression profiles that are neither too smooth nor too rough. Selecting suitable hyperparameters for the parameter priors is important to avoid unrealistic regions of parameter space. We have found an empirical Bayes approach useful in this regard.

We need to estimate how much of the variation in the data is due to noise and how much is due to temporal variation in the underlying expression profiles. We use a simple approach that might slightly overestimate both sources of variation but works well enough in our experience. First we group the expression measurements by gene and capture time to calculate means and variances.

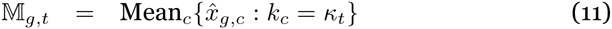

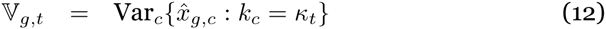

We estimate the gene-specific noise levels by assuming that all the within-time variation in the data is due to noise

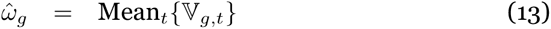

for each gene *g*. This ignores any effect the temporal variation may have had on the observed variance in the data and should overestimate the *ω*_*g*_.

To estimate the temporal variation, we examine the between time variation in the data. However we need to account for the effect of the covariance function Σ_*τ*_, which models the variation over time. We evaluate Σ_*τ*_ at the observed capture times to estimate the covariance of samples from the capture times. This gives a covariance matrix 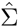 where

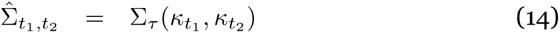

It can be shown (see Appendix 5.1) that a sample from a zero mean Gaussian with covariance matrix Σ is expected to have a variance of

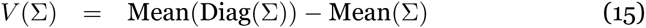

Using the linearity of this result the expected variance of samples from expression profile *y*_*g*_ at the capture times *κ*_1_,…,*κ*_*T*_ are expected to have variance equal to

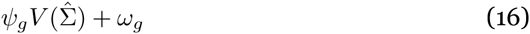

We slightly overestimate *ψ*_*g*_ by setting

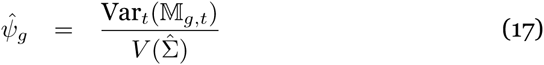

where the overestimation is caused by ignoring the effect of *ω*_*g*_ on the expected variance. This should be a small effect as we average over multiple cells per capture time.

Using these estimates for *ψ*_*g*_ and *ω*_*g*_ we empirically set the hyperparameters of the model as

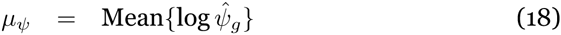

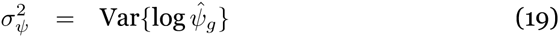

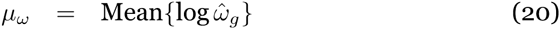

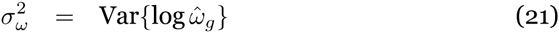

### 2.5 Inference

Our model is coded using the Stan probabilistic modelling language [27]. The Stan package provides various inference algorithms. In this work we have used the No-U-Turn Hamiltonian Markov chain Monte Carlo sampler (NUTS) [28]. In theory using a MCMC sampler gives us samples from the full posterior of the model. However the model is multimodal with respect to the pseudotime assignments and this makes it sometimes difficult for the sampler to mix samples from the full posterior. The multimodality occurs as there may be many pseudotemporal orderings of the cells that give smooth expression profiles. Moving between these modes is difficult for the sampler since in order to change the order of cells they must pass each other in pseudotime. If the cells’ expression profiles are sufficiently different the likelihood of the sampler passing this configuration can be very low. In these cases the sampler may only visit a few modes of the posterior. This difficulty in mixing is not unique to our model. Many other models such as K-means clustering exhibit similar behaviour. In these models it is common practice to use a single sample as a point estimate of the latent variables. Typically the sample with the highest probability under the model is selected. The Stan NUTS sampler provides *Ř* statistics that give confidence in the mixing over pseudotime [29]. These statistics can be evaluated on a dataset-by-dataset basis and a point estimate or the full posterior can be used for further analysis.

In order to further mitigate the pseudotime mixing problem we use a naive heuristic to initialise our MCMC chains. We sample many (by default, 6000) sets of pseudotimes from the prior and evaluate their likelihood (in combination with empirical Bayes estimates of the other parameters). We initialise our chains with the pseudotimes with highest likelihoods. This naive approach could almost certainly be improved upon but it is superior to using random samples from the prior to initialise the chains (data not shown).

### 2.6 Validation

In the results section we analyse specific data sets and validate the inferences from our model in several biological contexts. However we also wished to validate our model technically. We base this technical validation on the smoothness of expression profiles induced on held out genes. The held out genes are not used during model fitting and are only used in the validation stage. To evaluate the smoothness we developed a basic statistic to capture this concept. Given expression values 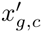 for a held out gene *g* over cells 1 ≤ *c* ≤ *C*, pseudotimes *τ*_1_,…, *τ*_*C*_ and an ordering *z*_1_,…,*z*_*C*_ such that

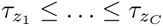

we define the roughness of the gene in terms of the differences of consecutive expression measurements under the ordering given by the pseudotimes

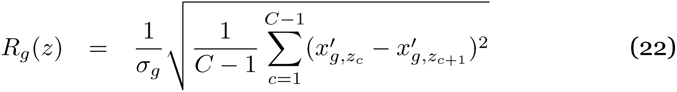

where *σ*_*g*_ is the standard deviation of the expression measurements. Clearly low *R*_*g*_ values should correlate with smooth profiles and high *R*_*g*_ values should correlate with rough profiles.

One benefit of defining *R*_*g*_ in terms of the pseudotime ordering rather than the pseudotime itself is that it is easy to generate random orderings under a suitable null hypothesis. The null hypothesis we use is that the cells are ordered by capture time but within a capture time are equally likely to have any order. That is, we generate random orderings that respect the capture times. We use a one-sided t-test to determine if the mean of the roughness of the pseudotime orderings is less than the mean of the roughness of orderings drawn under the null hypothesis. Defining *R*_*g*_ in terms of the ordering rather than the actual pseudotime also allows us to use it to compare the roughness of orderings from other methods such as Monocle.

### 2.7 Availability

The model described in this paper is available as an R package on github (https://github.com/JohnReid/DeLorean/). The code is open source and is available under a liberal MIT license. Code to replicate the results in this paper is available as vignettes in the package and the necessary data is also provided with the package.

## 3 Results

We used our model to analyse three sets of data from three different organisms assayed using three different technologies: whole leaf *Arabidopsis thaliana* microarrays [30]; nCounter single cell profiling of a human prostate cancer cell line [6]; and single cell RNA-seq of mouse dendritic cells [11].

### 3.1 The response of *Arabidopsis* to infection

Windram et al. examined the response of *Arabidopsis thaliana* to infection by the necrotrophic fungal pathogen *Botrytis cinerea* [30]. They generated highresolution time series over 48 hours for an infected condition and a control condition. We investigated if our model could estimate the correct order for the samples if their exact capture times were withheld.

#### 3.1.1 The model correctly estimates withheld sample times

Windram et al. measured expression levels every two hours resulting in 24 distinct capture time points. We grouped these 24 time points into four low-resolution groups, each consisting of six consecutive time points. We then asked our model to estimate the pseudotimes associated with each sample but only provided it with the low-resolution group labels. Our model is not suitable for fitting thousands of genes simultaneously so we fit 150 of the genes mentioned in the text of Windram et al.’s publication [30].

Our model estimated pseudotimes for each sample in the infected condition (see Figure 1). In general the mixing of the posterior was good as quantified by the 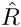 statistics (all but two of the *τ*_*c*_ 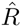 statistics were less than 1.15). The profiles induced by the inferred pseudotimes were smooth (see Figure 2).

**Figure 1:**
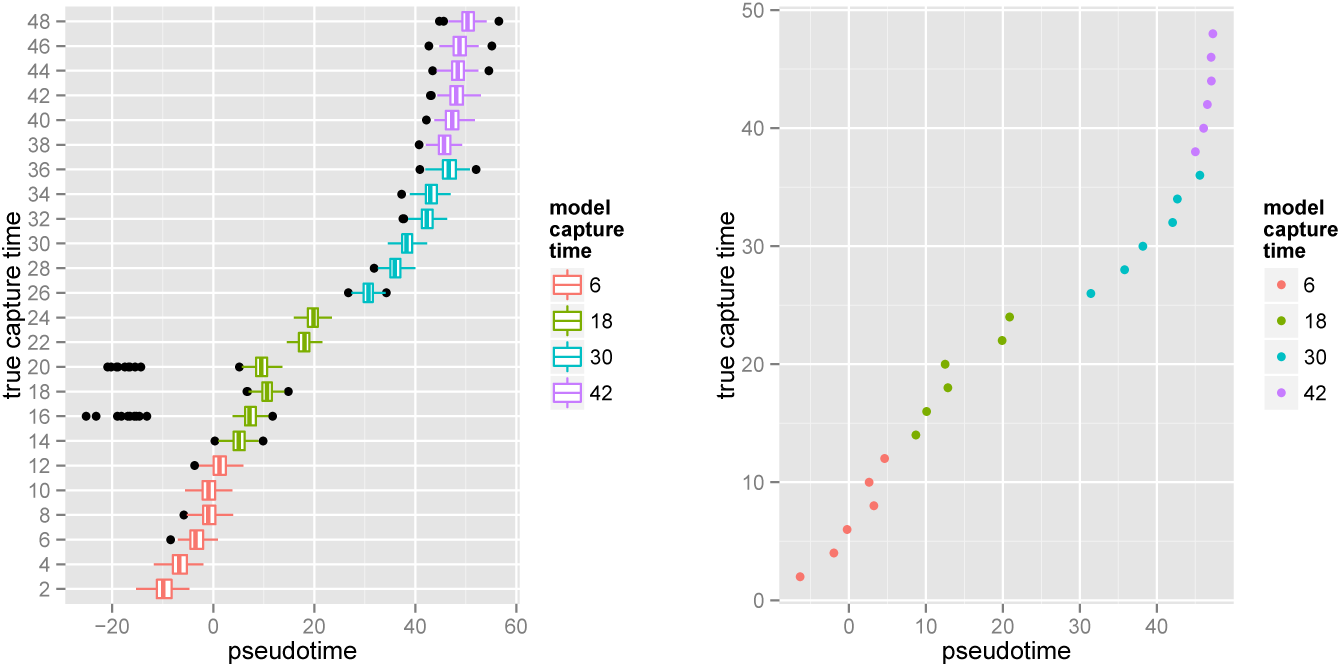
Pseudotime estimates for the samples from Windram et al.’s *Arabidopsis* data. *Left*: Boxplots of the full pseudotime posteriors. The estimated pseudotimes are in good agreement with the true capture times. The model tends to spread the samples out around the 20 hour mark in pseudotime. Presumably the expression profiles vary the most at this point. Also the samples are spread out more broadly in pseudotime (between −20 and 60 hours) compared to the true capture times. *Right*: The pseudotimes estimated by the best sample from the posterior plotted against the true capture times.

The Spearman correlation between estimated pseudotimes from the posterior and the true capture times was high (posterior mean *ρ* = .993) (see Figure 3 top left). The correlation for the best posterior sample had Spearman correlation *ρ* = .997.

**Figure 2:**
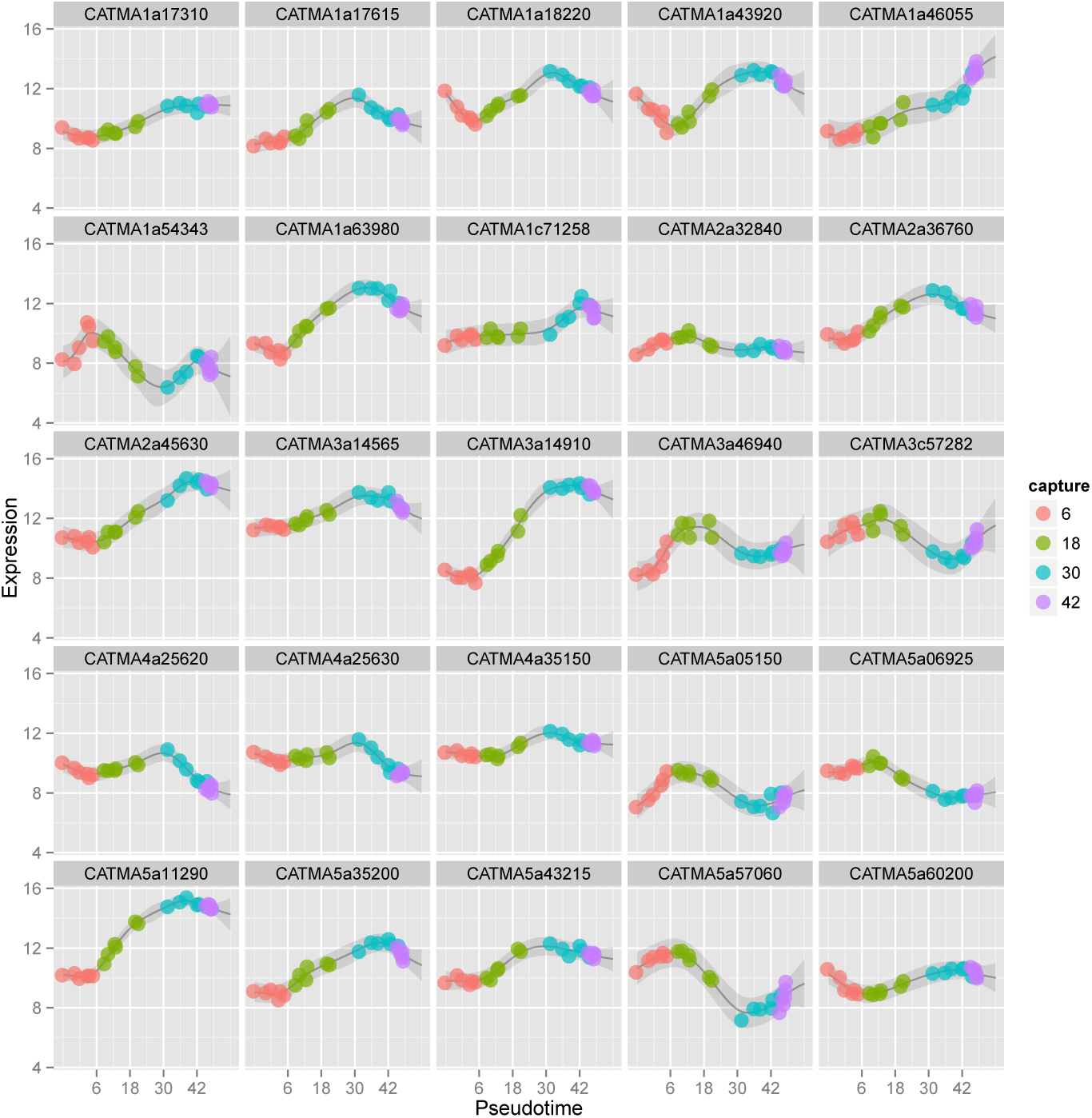
Expression profiles of selected genes over pseudotime. The expression data 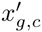 are shown as points coloured by their obfuscated capture time. The expected posterior mean of each profile is shown as a grey line and the shaded grey area in each profile represents the posterior uncertainty up to two standard deviations away from the mean.

#### 3.1.2 Our model fits the data better than Monocle

We also used the Monocle algorithm [12] to predict pseudotimes for the same 150 genes. Monocle was unable to recover the capture times for cells from the first low resolution group (see Figure 3). The Spearman correlation between Monocle’s estimated pseudotimes and the true capture times was not as high (*ρ* = 0.927) as that for our estimates. Monocle’s difficulty in resolving the correct ordering can be explained by its inability to use prior information that could resolve the first two groups of cells.

**Figure 3:**
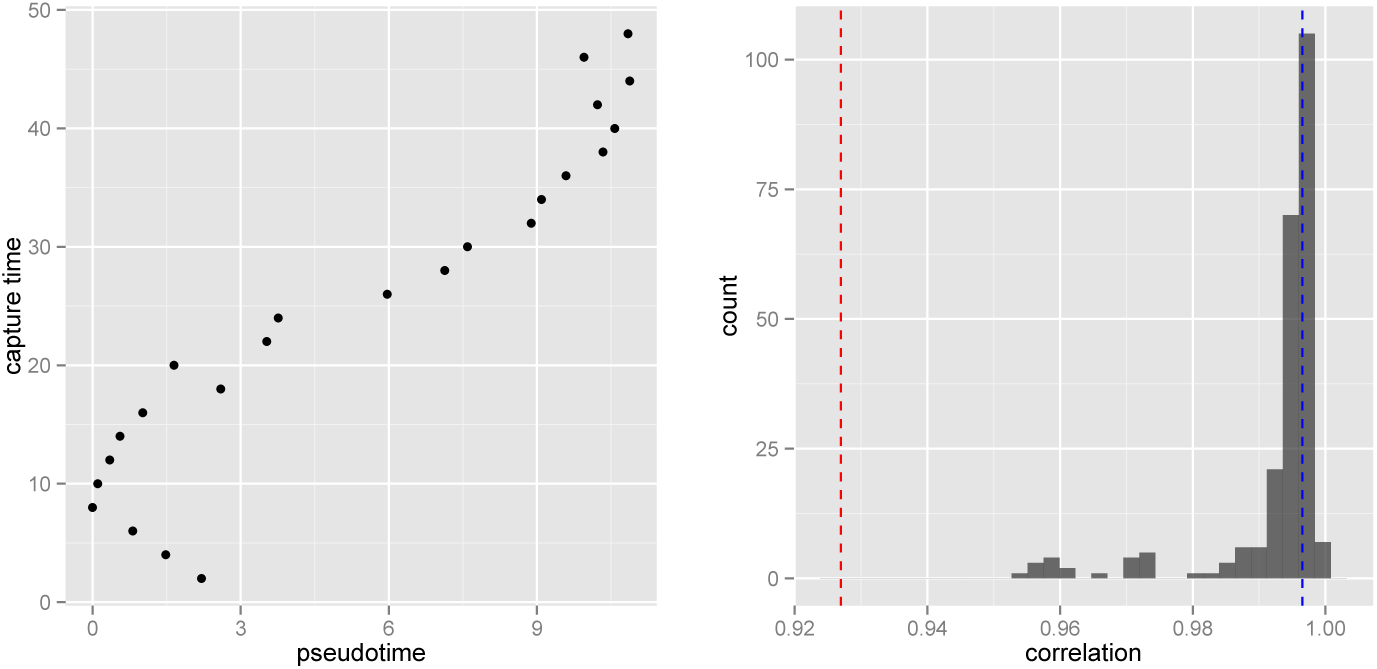
*Left*: Pseudotimes predicted by the Monocle algorithm (*ρ* = 0.927). *Right*: Posterior of the Spearman correlation between estimated pseudotimes from our model and true capture times. The Spearman correlation of the Monocle pseudotimes with the true capture times is shown as a red dashed line. The Spearman correlation of the best sample with the true capture times is shown as a blue dashed line.

#### 3.1.3 The pseudotimes induce smooth profiles on held out genes

We calculated roughness statistics *R*_*g*_ (see Section 2.6) for the 50 genes that we had not used to fit the model and averaged over genes. We did the same for 1000 pseudotime orderings sampled under the null hypothesis. The posterior mean of the *R*_*g*_ of the pseudotimes estimated by our model were significantly smaller than those from the null hypothesis (*p <* 10^−15^ one-sided t-test) (see Figure 4).

### 3.2 The effect of cell cycle on single cell gene expression

McDavid et al. were interested in the effect of the cell cycle on the single cell gene expression [6]. They assessed this effect by assaying the expression levels of 333 genes in 930 cells across three human cell lines using nCounter single cell profiling [31]. Based on these data they concluded that cell cycle explains just 5% to 17% of expression variability.

**Figure 4:**
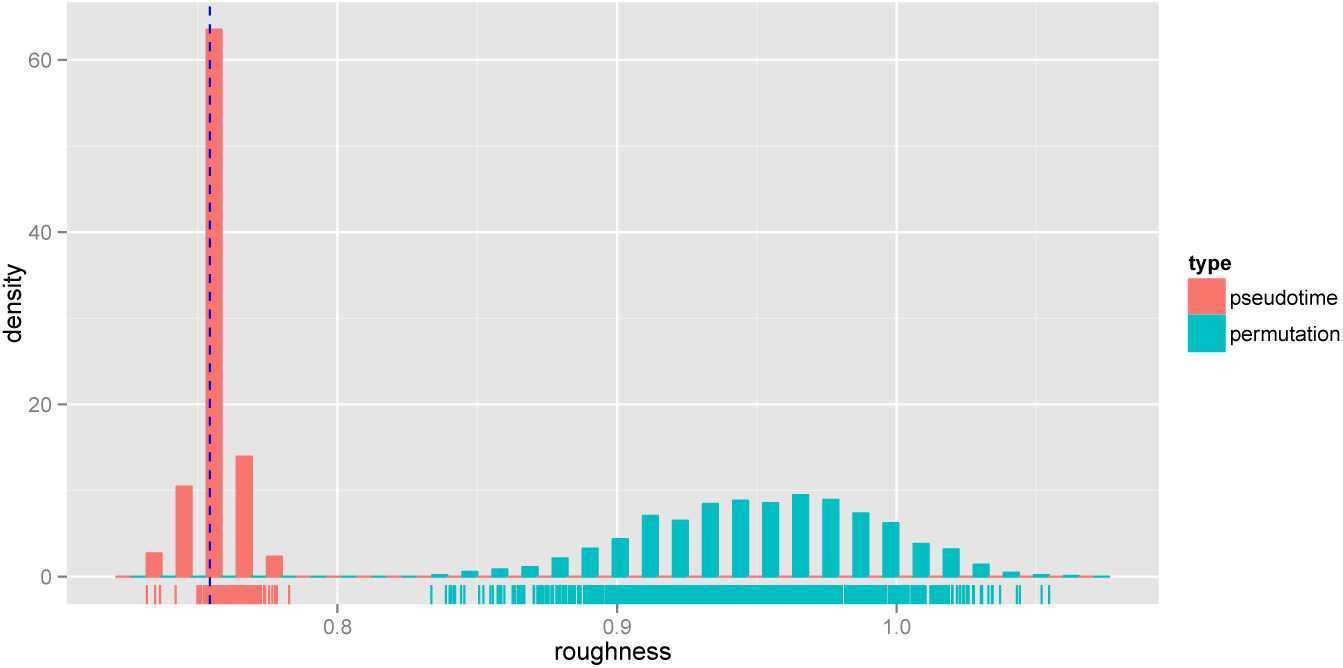
Roughnesses of samples from the posterior of our model (pink) and from draws from the null hypothesis (cyan). The roughness of the best sample from our posterior is shown as a dotted blue line.

CycleBase [32] is a database of cell cycle related genes and time series expression data. It contains metadata including the time in the cell cycle at which expression peaks for cell cycle related genes. To evaluate our model, we assessed how closely the peaks in the expression profiles estimated by our model from McDavid et al.’s data matched the CycleBase peak times. Additionally as a baseline, we compared peaks estimated from the raw expression data by a naive algorithm to the Cyclebase peak times.

#### 3.2.1 The model recovers cell cycle peak times

To fit our model we selected 37 cells at random from the PC3 human prostate cancer cell line and chose the top 56 differentially expressed genes according to McDavid et al.’s differential expression test. We mapped cells identified by McDavid et al. as G0/G1, S, and G2/M to capture times of 1, 2 and 3 respectively. We used a length scale of 5 and set 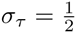. To model the cyclic nature of the cell cycle, we used a periodic covariance function with period Ω = 3. This ensured the expression profiles were periodic and transitions between all the cell cycle phases were consistent. Our model did not mix well with many 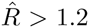 hence for further analysis we used the sample with highest log probability. We show expression profiles from this sample in Figure 5.

In order to test the fit of our model we estimated peak times from the expression profiles fit by the model and compared these to peak times as defined by the CycleBase database. To quantify this fit we calculated the root mean squared error (RMSE) between the CycleBase defined peak times and our estimates (RMSE=16.7).

We wished to understand how well our model estimated peak times compared to naive estimates. We made naive estimates from the raw expression data as follows. Each cell in McDavid et al.’s data had been labelled with one of the cell cycle phases. We identified the cell with maximal raw expression value for each gene. The middle of the cell cycle phase with which this cell was labelled was used as the naive estimate of the gene’s peak time. These estimated peak times had a RMSE of 21.3 which is 28% larger than the RMSE of our estimated peak times. This demonstrates that the our model’s expression profiles capture information present in the data at a higher temporal resolution than the raw labels.

**Figure 5:**
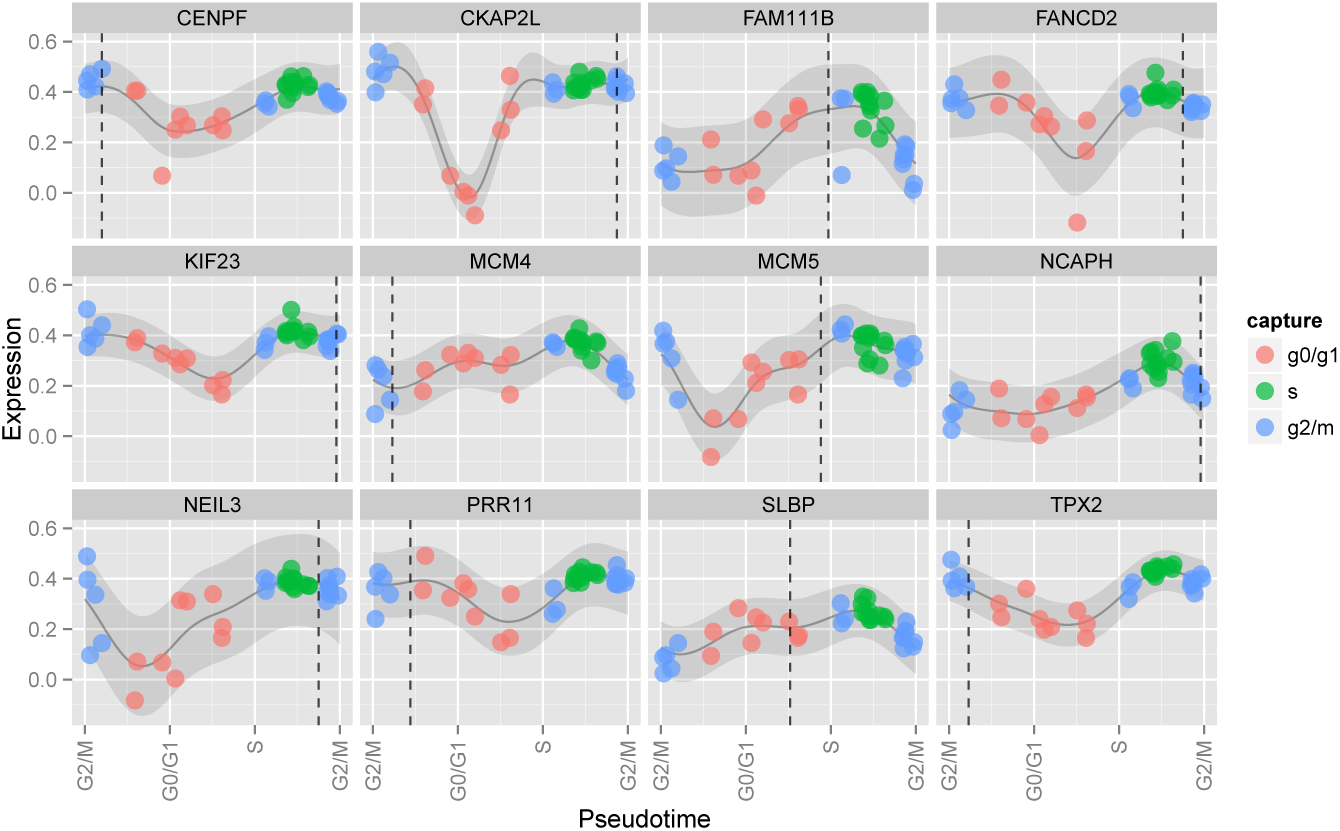
Expression profiles over pseudotime from the McDavid et al. cell cycle data. The pseudotimes are those from the best sample. Note the circular *x* axis: the first and last labels are both for the G2/M stage. The genes were selected based on high ratios of temporal variance to noise. Each point represents the expression of the given gene in a cell. The points are coloured by the cell cycle stage with which the cell was labelled by McDavid et al. The dark grey line represents the posterior mean of the expression profile for the gene and the shaded grey ribbon represents two standard deviations either side of this mean. The vertical dotted lines are the peak times as defined by the CycleBase database.

### 3.3 Paracrine signalling in mouse dendritic cells

Shalek et al. generated cross-sectional time courses of the response of primary mouse bone-marrow-derived dendritic cells in three separate conditions using single-cell RNA-seq [11]. We analysed the data on the lipopolysaccharide stimulated (LPS) condition using our model.

#### 3.3.1 The model identifies precocious cells

Shalek et al. identified a core antiviral module of genes that are expressed in conditions such as LPS after two to four hours. They also identified two cells captured at one hour that had this module switched on precociously. Other cells captured at one hour did not express the genes in this module. This concept that some cells can progress through pseudotime faster than others is exactly the concept that our model is designed to capture. We were interested to establish if our model could place these cells at later pseudotimes than other cells captured at one hour.

To fit our model we sampled 37 cells from the LPS condition including the two precocious cells captured at one hour. Shalek et al. defined several gene modules in their publication that show different temporal patterns of expression across the LPS time course. We selected the 74 genes from the clusters Id, IIIb, IIIc, IIId with the highest temporal variance relative to their noise levels. We set *σ*_*τ*_ = 1 and used a length scale of 5. The model mixed well with all but four of the cells’ pseudotimes having a 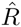 of less than 1.2.

Figure 6 shows the module scores of the core antiviral genes (as defined by Shalek et al.) over pseudotime. The two precocious cells have been fit with a pseudotime in the middle of the two hour capture cells.

**Figure 6:**
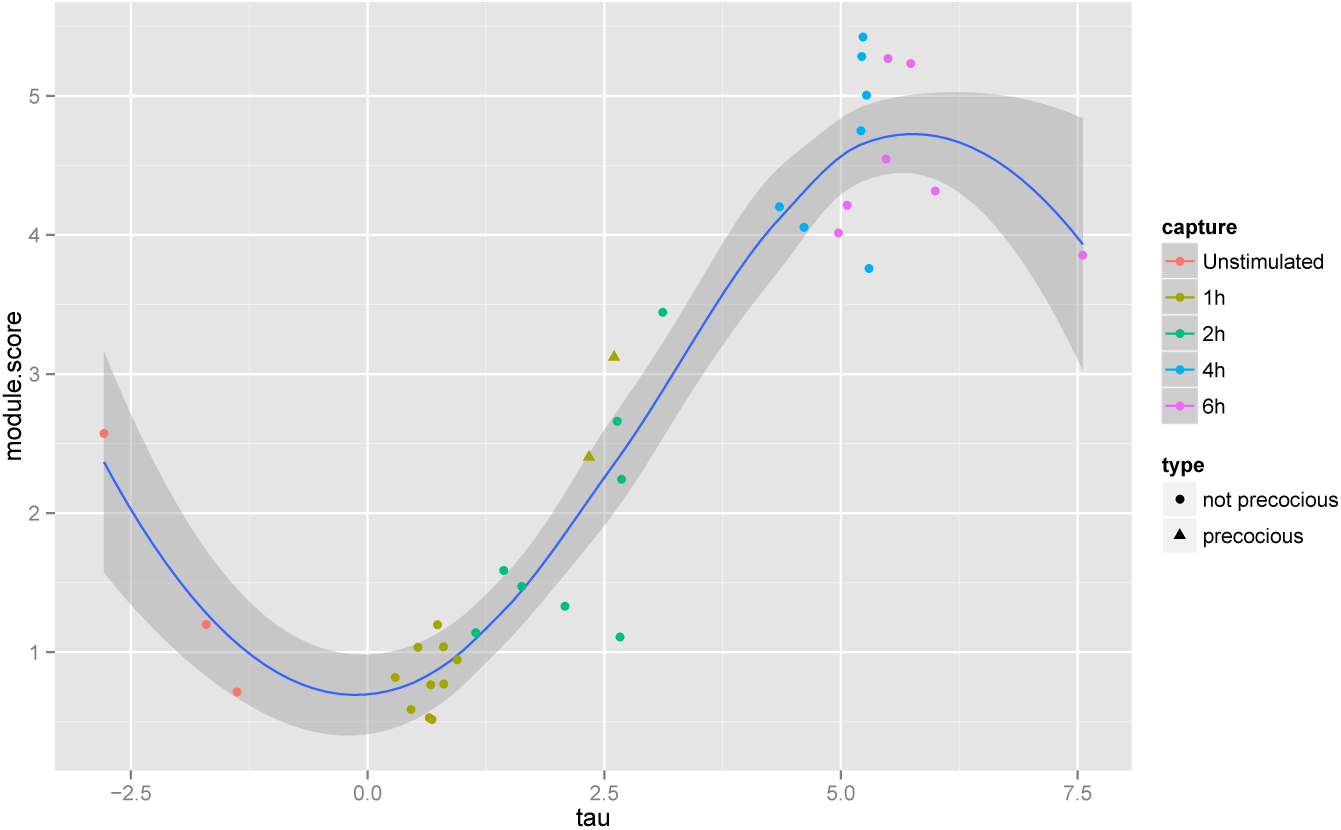
The module score (as defined by Shalek et al.) of core antiviral genes over pseudotime. The two precocious cells captured at one hour are plotted as triangles. These two cells have been placed at a later pseudotime than the other cells captured at one hour. A Loess curve has also been plotted through the data.

#### 3.3.2 The model identifies smooth expression profiles

We calculated roughness statistics *R*_*g*_ (see Section 2.6) for 100 genes that we had not used to fit the model and averaged over genes. We did the same for 1000 pseudotime orderings sampled under the null hypothesis. The posterior mean of the *R*_*g*_ of the pseudotimes estimated by our model were significantly smaller than those from the null hypothesis (*p <* 10^−15^ one-sided t-test) (see Figure 7).

**Figure 7:**
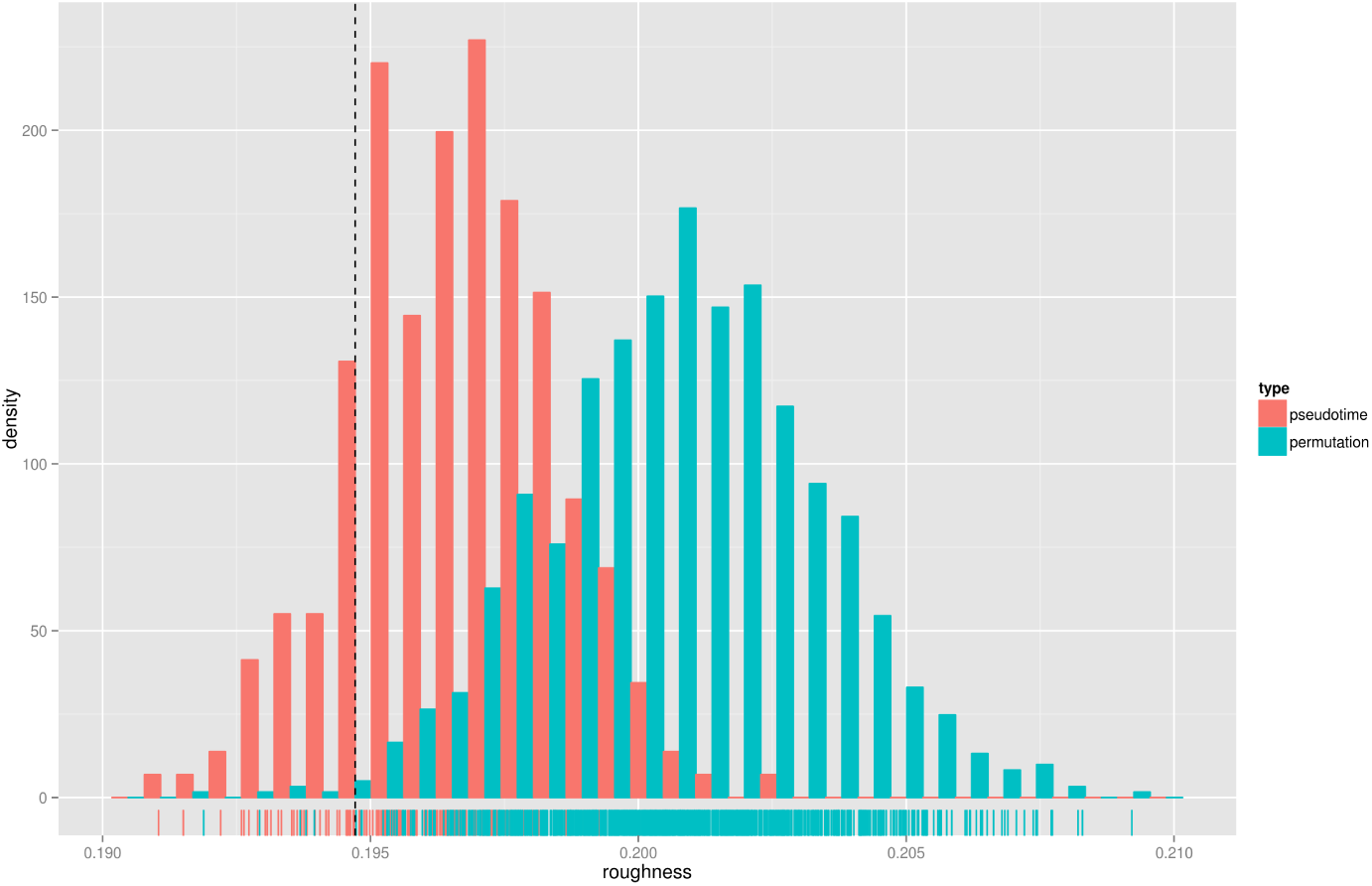
Roughnesses of samples from the posterior of our model (pink) and from draws from the null hypothesis (cyan). The roughness of the best sample from our posterior is shown as a dotted blue line.

## 4 Conclusions

We have presented a principled probabilistic model that accounts for uncertainty in the capture times of cross-sectional time series. We have fit our model to three separate data sets each using a different biological assay (microarrays, single cell nCounter and single cell RNA-seq) in three organisms (human, mouse and *Arabidopsis*). Our model provided plausible estimates of pseudotimes on all the data sets. We validated these estimates technically by evaluating the smoothness of the expression profiles of held out genes in two of the data sets. These profiles are significantly smoother than expected under the null model. In addition we validated the estimates biologically using obfuscated capture times (in the *Arabidopsis* data set), data from separate experiments (cell cycle peak times) and independent analyses (identification of precocious cells). Overall these results demonstrate that uncertainty in the temporal dimension should not be ignored in cross-sectional time series of single cell data and that our method captures and corrects for these effects.

Our method has a number of attractive attributes. It explicitly estimates pseudotimes in contrast to methods such as Monocle and Wanderlust which estimate orderings of cells. The pseudotimes are on the same scale as the experimental capture times. The orderings estimated by Monocle and Wanderlust have no scale. Additionally, the orderings estimated by Monocle and Wanderlust do not quantify the differences in expression between consecutive cells. Consecutive cells in the ordering could have similar or diverse expression profiles. In our model, consecutive cells that have diverse expression profiles are placed further apart in pseudotime than similar cells. Thus our pseudotime estimates quantify the rate of change of the system. For example, in the *Arabidopsis* example we analysed, the cells are spread out in pseudotime around the 20 hour mark (Figure 1) suggesting changes in expression levels in response to the infection are greatest at this time point.

Our method uses Gaussian processes which are a natural framework to model noisy expression profiles. Gaussian processes are well established probabilistic models for time series. They provide more than just point estimates of the profiles, they also provide a measure of posterior uncertainty. This is useful in downstream analyses such as regulatory network inference. A Gaussian process model is characterised by its covariance function and associated parameters and the covariance functions in our model have interpretable parameters: gene-specific temporal variation and noise. We have also demonstrated how a Gaussian process framework is suitable for modelling periodic expression profiles such as cell cycle expression profiles. The primary limitation of Gaussian processes for our model is that inference complexity scales cubically in the number of samples. For this reason our method is not applicable to data from many hundreds or thousands of cells like Monocle and Wanderlust.

Inference in our model is performed using Markov chain Monte Carlo. This technique provides a full posterior distribution over the model parameters. However mixing over the pseudotime parameters in our model can be difficult and we found that our model did not mix well when fit to the cell cycle data set. In this case, we analysed expression profiles from the sample with highest log probability and found they estimated cell cycle peak times well.

Single cell assays give us an exciting opportunity to explore heterogeneity in populations of cells. As the technology develops and the cost of undertaking such assays drops, they are destined to become commonplace. Also high throughput longitudinal studies remain impractical and for the foreseeable future the majority of such time series will be cross-sectional in nature. Until this changes there will be challenges associated with estimating uncertainty in the capture times and variation in the rate of progress of individual cells through a system. Our method explicitly models these effects and is a practical tool for analysis of such cross-sectional time series. Furthermore in contrast to Wanderlust our method only depends on open source software and is available under a liberal open source license.

## 5 Appendix

### 5.1 Variance of draw from multivariate Gaussian

Suppose we have a zero-mean Gaussian

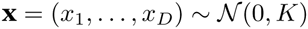

we wish to estimate the expected sample variance

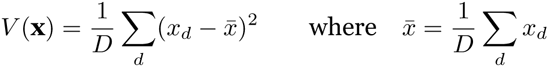

and all sums are from 1 to *D*. Setting 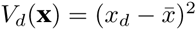 we have

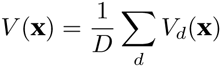

and

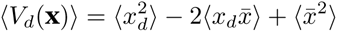

but

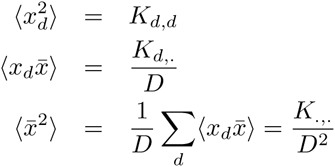

where a dot represents summation over that index, so

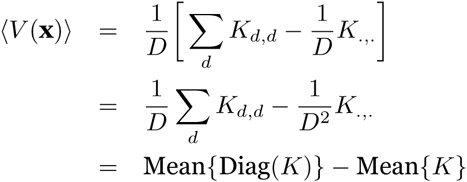

